# Cumulative effects of social stress on reward-guided actions and prefrontal cortical activity

**DOI:** 10.1101/817361

**Authors:** Florent Barthas, Melody Y. Hu, Michael J. Siniscalchi, Farhan Ali, Yann S. Mineur, Marina R. Picciotto, Alex C. Kwan

## Abstract

When exposed to chronic social stress, animals display behavioral changes that are relevant to depressive-like phenotypes. However, the cascading relationship between incremental stress exposure and neural dysfunctions over time remains incompletely understood. Here we characterize the longitudinal effect of social defeat on goal-directed actions and prefrontal cortical activity in mice, using a head-fixed sucrose preference task and two-photon calcium imaging. Behaviorally, stress-induced loss of reward sensitivity intensifies over days. Motivational anhedonia, the failure to translate positive reinforcements into future actions, requires multiple sessions of stress exposure to become fully established. For neural activity, individual layer 2/3 pyramidal neurons in the Cg1 and M2 subregions of the medial prefrontal cortex have heterogeneous responses to stress. Changes in ensemble activity differ significantly between susceptible and resilient animals after the first defeat session, and continue to diverge following successive stress episodes before reaching persistent abnormal levels. Collectively, these results demonstrate that the cumulative impact of an ethologically relevant stress can be observed at the level of cellular activity of individual prefrontal neurons. The distinct neural responses associated with resilience versus susceptibility raises the hypothesis that the negative impact of social stress is neutralized in resilient animals, in part through an adaptive reorganization of prefrontal cortical activity.

## Introduction

Stress is associated with increased risk for multiple neuropsychiatric disorders, including major depression (1). A prevailing framework for stress-related disorders is the idea of allostatic load, which posits that although acute stress may promote adaptation, repeated exposures over a prolonged period of time can lead to persistently heightened reactions and maladaptations (2, 3). Accordingly, there is growing evidence that acute and chronic stress are associated with distinct alterations in the mammalian brain (4, 5). However, effects of stress on behavior and neural functions are generally sampled at the beginning and at the end of chronic stress exposure, whereas the adaptations across successive stress episodes is not understood. One difficulty is that individuals may exhibit different degrees of resilience, and therefore their time-dependent responses to repeated stress may be variable. Another difficulty is that longitudinal measurements are needed to track the dynamic relationship. For example, depressive-like phenotypes in rodents are thought to arise only after chronic stress, yet the timing of when the dysfunction is precipitated remains poorly understood.

In this study, we characterize the time course of stress effects on motivated behavior and prefrontal cortical activity in mice. Motivated behavior – specifically the capacity to select appropriate actions based on reinforcement feedback – is an essential aspect of learning. Deficits in reward-guided behavior can be quantified precisely using instrumental decision-making tasks in multiple animal species and therefore have translational significance (6). For example, in humans, stress blunts goal-directed actions and promotes habitual responses (7). In agreement, rodents subjected to chronic stress paradigms exhibit a diminished ability to modify actions flexibly in an outcome-dependent manner (8, 9).

A classic assay to measure reward-guided behavior in stressed rodents is sucrose preference. In this assay, an animal is presented with two bottles, one containing water and the other sucrose solution, and is given a certain duration of time to consume the fluids. A lack of preference for the sucrose solution, which is normally favored by rodents, is considered to be an indicator of anhedonia. Prior studies have reported reduced sucrose preference following chronic stress, such as after repeated social defeat (10, 11). However, reduced preference can arise for at least two reasons. First, animals could be impaired in their ability to sense or gain pleasure from sucrose. This would relate to the emotional reactivity to the reward. Second, the sucrose may remain desirable, but animals could fail to translate that positive reinforcement into future actions. These two components of reward-guided behavior are well-known, and the corresponding deficits have been termed ‘appetitive anhedonia’ and ‘motivational anhedonia’ (12, 13). Unfortunately, the conventional sucrose preference metric does not dissociate these two forms of reward processing deficits (14), so a more sophisticated paradigm is necessary to understand the underlying mechanisms of stress-induced changes in reward-guided behavior.

Reward-guided behavior is thought to involve the distributed cortical-basal ganglia network including the prefrontal cortex (6). Consistent with the observed deficits in reward processing, the effects of chronic stress on the prefrontal cortex can be detected at many levels, including the morphological, molecular, and cellular (4, 5). For example, in rodents, prolonged stress exposure causes structural atrophy, including dendritic retraction and synapse loss (8, 15, 16), and disrupts synaptic plasticity and synaptic transmission in the medial prefrontal cortex (17, 18). Although it is obvious that the structural and synaptic alterations must compromise prefrontal cortical function, details of stress effects on neural activity *in vivo* are less clear. There have been several studies of stress effects on cortical activity patterns (19-22), but they provide few details on the effect of escalating stress burden over time.

Given the gaps in our current knowledge, the main goal of this study is to determine how social stress influences reward-guided behavior and prefrontal cortical activity over time. To this end, we have designed a self-paced, instrumental sucrose preference task for the head-fixed mouse. Detailed characterization of the lick microstructure and a computational model of the behavior allows us to attribute behavioral alterations more confidently to motivational or appetitive anhedonia. To characterize the concomitant changes in prefrontal cortical activity, we used two-photon calcium imaging to track neural ensembles over two weeks and visualize spontaneous activity. Using these approaches, we sampled behavioral performance and neural activity longitudinally before, during, and after successive social defeat episodes, for susceptible and resilient animals. The results reveal a progressive emergence of motivational anhedonia and cortical activity maladaptations as a function of cumulative social stress and resilience.

## Results

### A self-paced, instrumental sucrose preference task for the head-fixed mouse

We designed a self-paced, instrumental sucrose preference task to characterize reward-directed actions. We had three major criteria for the task: Actions are self-paced to engage voluntary behavior; Sucrose reinforcements are varied to manipulate the motivational level; Animal is head-fixed to enable cellular-resolution optical imaging, but mobile to minimize stress during behavioral testing. To meet these criteria, we incorporated concepts from prior studies involving ratio schedules (23), incentive contrast (24, 25), and head fixation for awake, mobile mice (26). During the task, the head-fixed mouse is able to walk atop a rolling Styrofoam ball or a rotating disk (**Fig. 1a**). Tongue licks on a spout are detected by an electronic touch detector circuit and reinforced on a tandem fixed-ratio 10, fixed-interval 5 schedule (FR10-FI5; **Fig. 1b**). Specifically, animal has to first complete an FR10 schedule, for which the mouse must lick 10 times to receive ∼4 µL of fluid, and this is immediately followed by an FI5 schedule, which is effectively a 5 s timeout after the reinforcement to avoid counting consummatory licks towards the subsequent FR10 schedule. The spout is constructed by fusing three needle tips. Each tip can independently deliver water, 3% sucrose, or 10% sucrose solutions. For simplicity, we will refer to water as 0% sucrose solution. We chose 10% as the highest concentration, because it promotes maximal consumption in C57BL/6J mice (27). The task employs a block design (**Fig. 1c**). Each block lasts 60 s and is associated with one fluid type. The mouse can initiate a block by completing an FR10 schedule, and then has 60 s to complete more FR10 schedules to obtain the same type of reinforcement. The fluid type changes from block to block, always going from 0% to 3% to 10% to 3%, and then the sequence repeats again. Cycling through the fluid types allows the study of relative reward values, because a 3%-sucrose block preceded by a 0%-block has different successive incentive contrast compared to a 3%-sucrose block preceded by a 10%-block. We referred to these two kinds of 3%-blocks as 3% positive contrast (3%pc) and 3% negative contrast (3%nc) blocks. Overall, the self-paced, instrumental sucrose preference task allows us to measure how an animal adjusts its actions in response to differences in absolute reward values (i.e., 0% vs. 10%) and relative reward values (i.e., 3%nc and 3%pc).

**Figure 1:**
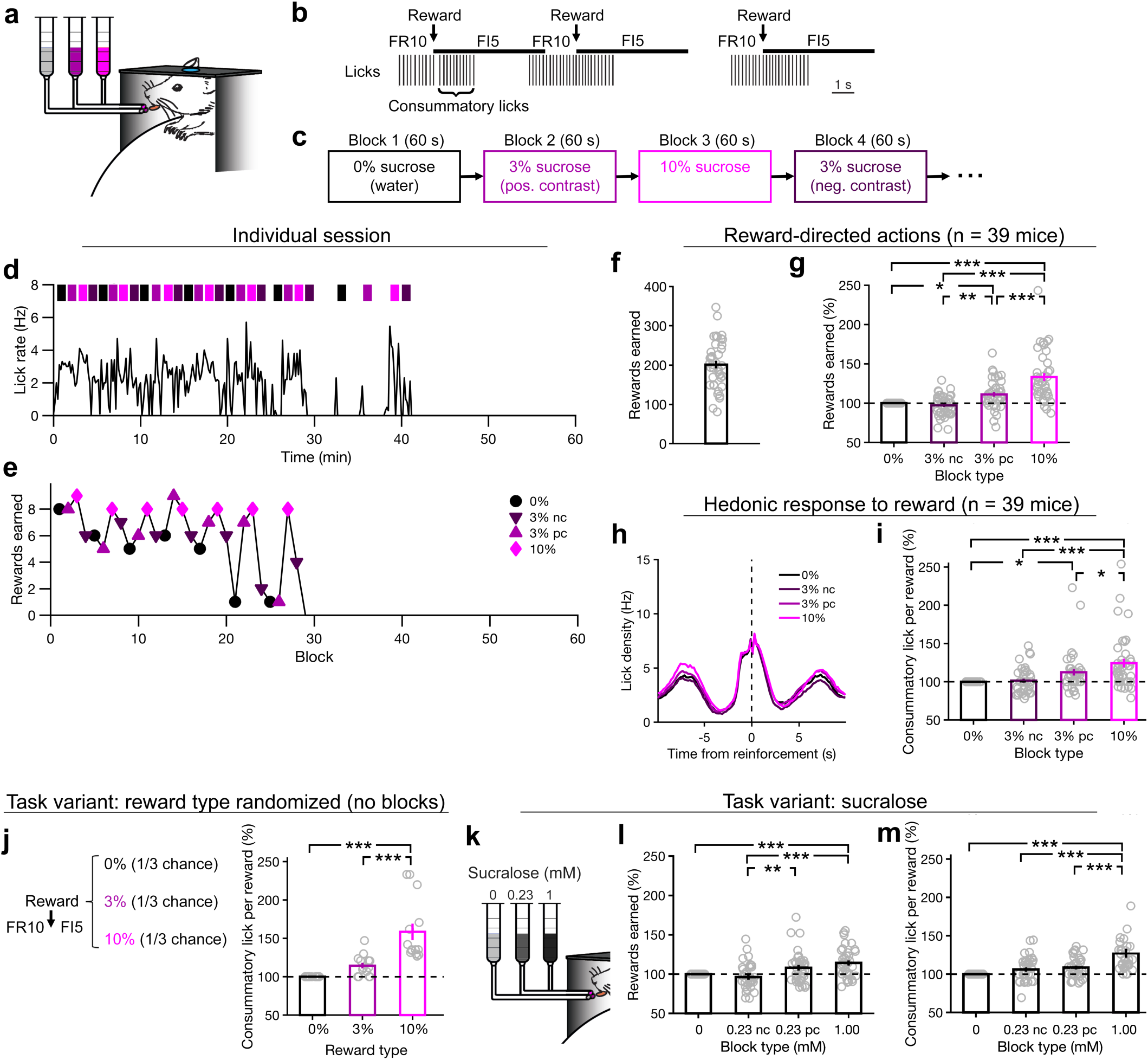
A self-paced, instrumental sucrose preference task for the head-fixed mouse. **(a)** Schematic of the experimental setup. The head-fixed mouse had access to a composite lick spout with three openings, each of which could deliver fluid independently. The mouse sat atop a skewered sphere or a treadmill and could run forwards or backwards. **(b)** The task was based on a tandem fixed-ratio (FR), fixed-interval (FI) schedule. To illustrate the FR10-FI5 schedule, licks detected and reinforcement delivered from an actual behavioral session were plotted. **(c)** The task had a block design, cycling through multiple types of reinforcement including 0%, 3%, and 10% sucrose solutions. The sequence was always 0%, 3%, 10%, 3%, and then 0%, 3%, 10%, 3% again, and so on. **(d)** Lick rate for a typical session. Lick rate was calculated in 10-s bins. The colored patches indicated the reinforcement type and duration of the blocks. Note that there were time gaps between the blocks, because a new block would begin only after the completion of a FR10 action sequence. **(e)** Rewards earned for each block. This is the same session as (d). **(f)** Rewards earned per session. Open circles, individual sessions; bar, mean ± s.e.m. **(g)** Rewards earned for each reinforcement type, normalized by rewards earned in 0% blocks, per session. Open circles, individual sessions; bar, mean ± s.e.m; Main effect of reinforcement type (F_3,114_ = 47.8; *P* = 4 × 10^−20^), comparisons: 0% vs. 3% pc, *P* = 0.012; 0% vs. 10%, *P* = 4 × 10^−9^; 3% nc vs. 3% pc, *P* = 0.0011; 3% nc vs. 10%, *P* = 4 × 10^−9^; 3% pc vs. 10%, *P* = 1 × 10^−7^, ANOVA with post hoc Tukey-Kramer test. **(h)** Mean lick density relative to the time of reinforcement, plotted separately for each reinforcement type. **(i)** Consummatory licks per reward for each reinforcement type, normalized by the number in 0% blocks. Open circles, individual sessions; bar, mean ± s.e.m; Main effect of reinforcement type (F_3,114_ = 9.0; *P* = 6 × 10^−8^), comparisons: 0% vs. 3% pc, *P* = 0.02; 0% vs. 10%, *P* = 5 × 10^−7^; 3% nc vs. 10%, *P* = 2 × 10^−6^; 3% pc vs. 10%, *P* = 0.03, ANOVA with post hoc Tukey-Kramer test. **(j)** A separate cohort of mice was trained on a variant of the task with the same FR10-FI5 schedule, however reinforcement type was randomized for each reward. Open circles, individual sessions; bar, mean ± s.e.m; Main effect of reinforcement type (F_2,26_ = 25.0; *P* = 9 × 10^−7^), comparisons: 0% vs. 10%, *P* = 8 × 10^−7^; 3% vs. 10%, *P* = 1 × 10^−4^, ANOVA with post hoc Tukey-Kramer test. **(k)** Another separate cohort of mice was trained on a variant of task in which the reinforcement types were 0, 0.23, and 1 mM sucralose. **(l)** Same as **(h)** except for sucralose task. Main effect of reward type (F_3,105_ = 10.6; *P* = 4 × 10^−6^), comparisons: 0 vs. 1.00, *P* = 0.0006; 0.23 nc vs. 0.23 pc, *P* = 0.006; 0.23 nc vs. 1.00, *P* = 8 × 10^−6^, ANOVA with post hoc Tukey-Kramer test. **(m)** Same as **(i)** except for sucralose task. Main effect of reward type (F_3,105_ = 15.9; *P* = 1 × 10^−8^), comparisons: 0 vs. 1.00, *P* = 2 × 10^−8^; 0.23 nc vs. 1.00, *P* = 9 × 10^−6^; 0.23 pc vs. 1.00, *P* = 1 × 10^−4^, ANOVA with post hoc Tukey-Kramer test. Sample sizes were 39 sessions from 39 mice **(f, g, h, i)**, 14 sessions from 2 mice **(j)**, and 39 sessions from 9 mice **(k, l, m)**. * *P* < 0.05, ** *P* < 0.01, *** *P* < 0.001. In the list above, if a potential post hoc comparison was unnoted, *P* > 0.1.

### Mice are sensitive to sucrose concentration and incentive contrast

To encourage animals to perform the self-paced, instrumental sucrose preference task, mice were fluid-restricted. On alternate days, animals would either obtain its entire fluid intake from the task, or be given 15 minutes of *ad libitum* access to water. **Figure 1d** shows a typical behavioral session for an adult male C57BL/6J mouse. The animal responded for about 30 min before stopping due to satiation. Plotting the number of rewards earned on a block-by-block basis revealed a sawtooth-like modulation, which matched the rises and falls of the sucrose concentration (**Fig. 1e**) As expected, the mouse responded most vigorously and earned more rewards for the 10%-blocks. Prior study has shown that mice suppress intake when they anticipate more palatable fluids in the near future (28). This can be seen in **Figure 1e**, because the mouse ceased licking for the duration of a few 60-s blocks (i.e., blocks 21, 25, and 26 with 1 reward each, which is the minimum needed to initiate the block), and then resumed when a more palatable fluid became available.

We analyzed performance from 39 sessions (n = 39 mice). On average, animals earned 201 ± 10 reinforcements per session (mean, s.e.m.; **Fig. 1f**). Mice were motivated to complete more FR10 schedule to obtain more reinforcements during blocks with more palatable fluids (*P* = 4 × 10^−20^, main effect of block type, one-way ANOVA; for details on the statistical analyses, refer to figure captions; **Fig. 1g**). In particular, animals obtained 33 ± 5% more rewards during 10%-blocks relative to 0%-blocks (*P* = 4 × 10^−9^, post hoc Tukey-Kramer test). Animals also responded significantly more during 3%pc-blocks than 3%nc-blocks (*P* = 0.0011, post hoc Tukey-Kramer test). Therefore, both the absolute and relative differences in reward values were relevant for performance on this task. Because ratio schedule is a well-known approach to induce goal-directed actions (23), these results emphasize the motivational component of action.

Rodents lick in bouts with characteristic frequencies and pauses. A ‘lick bout’ can be defined as a train of licks occurring less than 0.5 s apart (29). Indeed, during the self-paced, instrumental sucrose preference task, mice licked in bouts with a pause between successive reinforcements (**Fig. 1h**). A typical bout would therefore include 10 licks to complete the fixed-ratio schedule, and then additional consummatory licks after reinforcement (11.0 ± 0.5 licks for water; **Fig. 1b**). Animals had more consummatory licks as the concentration of sucrose was increased (*P* = 6 × 10^−8^, main effect of block type, one-way ANOVA; **Fig. 1i**). There is an extensive literature indicating that the number of consummatory licks is directly related to the emotional reaction to reinforcement (29, 30), and thus this measurement focuses on the appetitive component of action.

Basing our interpretation on prior literature, we have suggested that the numbers of rewards earned reflects the motivational component of the response, whereas consummatory licks reflect the appetitive component of the response. One way to test this dissociation explicitly is to try a task variant with no motivational component. To this end, animals were tested on a version of the task in which reinforcements were assigned randomly on a trial-by-trial basis, such that there is no motivation for the animal to adjust its instrumental actions because the reward value is unpredictable. For animals that have only been exposed to this task variant (n = 14 sessions, 2 mice), we found that the consummatory licks remained sensitive to the fluid type (*P* = 9 × 10^−7^, main effect of reward type, one-way ANOVA; **Fig. 1j**). Furthermore, sucrose is not only palatable, but also contains calories which may appeal to fluid-restricted animals. To exclude calorie as the factor underlying performance, we replaced sucrose with sucralose, which is a zero-calorie artificial sweetener. With 0, 0.23, and 1 mM sucralose solutions as the reinforcement types, mice remained motivated to earn more rewards and had more consummatory licks for increasing reward values (**Figs. 1k – m**).

### Motivational anhedonia in mice susceptible to social defeat stress

Although it is known that stress can induce anhedonia in rodents, the extent to which behavioral alterations can be attributed to motivational versus appetitive components, as well as the time course of the maladaptations, is not known. We therefore wanted to determine how repeated stress influences reward-directed actions. To induce stress, we used social defeat because of its ethological relevance for animals, and because of its construct and face validity (10). Moreover, we measured prefrontal cortical activity, and chronic social defeat stress is associated with maladaptations in the rodent frontal cortex (11, 21, 31). In these experiments (**Fig. 2a**), C57BL/6J mice received 10 days of social defeat involving encounters with aggressive resident CD-1 mice. Our procedures largely followed a standardized protocol (32), with one modification to decrease the intensity of each stress episode by reducing the duration of sensory interaction (33). Animals were tested on the self-paced, instrumental sucrose preference task at 2-day intervals before, during, and after the defeat sessions. In addition, to ensure that the defeat protocol had the expected effects on animals, mice were evaluated with a social interaction test on the day after the last defeat session.

**Figure 2:**
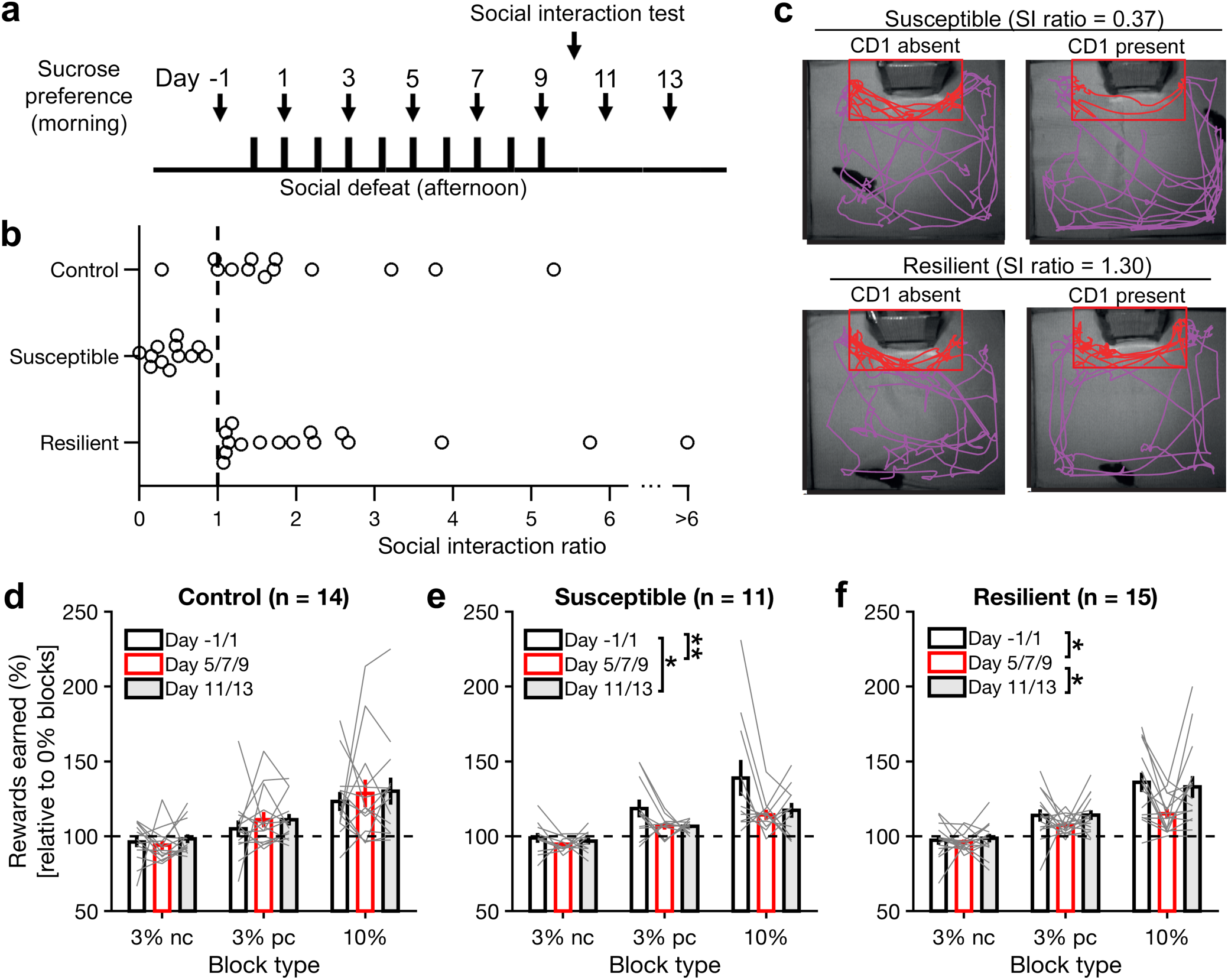
Motivational anhedonia as a sustained phenotype in susceptible, but not resilient, mice. **(a)** Timeline of the chronic social defeat stress and behavioral experiments. **(b)** Based on the social interaction ratio, mice subjected to social defeat were divided into resilient and susceptible individuals. Controls were handled but did not experience social defeat. Each circle in the beeswarm plot represents an animal. **(c)** Example movement trajectories for a susceptible and a resilient individual, when the CD1 aggressor mouse was present or absent in the interaction zone (red rectangle). **(d)** Rewards earned for each reinforcement type, normalized by rewards earned in 0% blocks, per session, for control animals. For each animal, the value was an average across days according to three stages: “pre” for day -1 and 1, “stress” for day 5, 7, and 9, and “post” for day 11 and 13. Gray line, individual animal; bar, mean ± s.e.m. Main effect of reinforcement type (F_2,117_ = 21.4, *P* = 1 × 10^−8^), but not for stage (F_2,117_ = 0.57, *P* = 0.6) or interaction (F_4,117_ = 0.16, *P* = 1), ANOVA with reinforcement type and stage as fixed factors, subject as random factor. **(e)** Same as **(d)** for susceptible animals. Main effect of reinforcement type (F_2,90_ = 20.0, *P* = 7 × 10^−8^) and stage (F_2,90_ = 6.16, *P* = 0.003), but not interaction (F_4,90_ = 1.28, *P* = 0.3), with comparisons: pre vs. stress, *P =* 0.005; pre vs. post *P =* 0.02; stress vs. post, *P =* 0.9, ANOVA with reinforcement type and stage as fixed factors, subject as random factor, and post hoc Tukey-Kramer test. **(f)** Same as **(d)** for resilient animals. Main effect of reinforcement type (F_2,126_ = 40.6, *P* = 3 × 10^−14^), stage (F_2,126_ = 5.05, *P* = 0.008), but not interaction (F_4,126_ = 1.70, *P* = 0.2), with comparisons: pre vs. stress, *P* = 0.013; pre vs. post, *P* = 1; stress vs. post, *P* = 0.02, ANOVA with reinforcement type and stage as fixed factors, subject as random factor, and post hoc Tukey-Kramer test. Sample sizes were 14 mice **(d)**, 11 mice **(e)**, and 15 mice **(f)**. * *P* < 0.05, ** *P* < 0.01, *** *P* < 0.001.

The cohort tested included 28 stressed and 14 control animals. During social interaction tests, an animal’s propensity for social interaction was characterized by calculating the social interaction ratio measured as time in the interaction zone with an aggressor divided by the time in the interaction zone without an aggressor. Based on this measure, defeated animals interacted less than control animals (stressed: 1.10, control: 1.60; *P* = 0.14, Wilcoxon’s rank-sum test). Furthermore, we used a unity threshold to divide defeated animals into susceptible and resilient subtypes, following the convention of earlier studies (10, 32). Among the stressed animals, 43% (12 out of 28) of the animals were classified as susceptible, versus 14% of the control mice if we had classified them using the same threshold (**Fig. 2b – c**). Collectively, these results confirm that the social defeat procedures had a relevant stress-dependent behavioral outcome.

We assessed the performance of control, susceptible, and resilient animals in the self-paced, instrumental sucrose preference task. We divided the behavioral data into pre-stress (day -1 and 1), during-stress (day 5, 7, and 9), and post-stress (day 11 and 13) stages. As expected, for control mice, rewards earned increased with the reward value, and this motivational effect was stable across days (*P* = 1 × 10^−8^, main effect of reward type; ANOVA with block type and stage as fixed factors and subject as random factor; **Fig. 2d**). By contrast, susceptible mice had diminished reward sensitivity that persisted into the post-stress stage (*P* = 7 × 10^−8^, main effect of reward type; *P* = 0.003, main effect of stage; **Fig. 2e**). This stress-induced loss of reward responsiveness was significant for both absolute and relative differences in reward values. Resilient mice also had a loss of reward sensitivity, however the decrease was transient during stress, as the sucrose preference quickly returned to pre-stress level following the last defeat session (*P* = 3 × 10^−14^, main effect of reward type; *P* = 0.008, main effect of stage; **Fig. 2f**). Therefore, animals subjected to repeated social defeat displayed motivational anhedonia, with the susceptible subtypes displaying a more prolonged phenotype.

### Delayed emergence of motivational anhedonia from cumulative effects of stress

Can we parameterize the factors that drove the motivational anhedonia? To answer this question, we turned to a model of self-paced actions based on motivational vigor. In the original model, an agent chooses an action out of multiple options by balancing the potential reward against the energetic costs for performing that action, following the principles of reinforcement learning (34). For reduced tasks with only one type of action, the model simplifies substantially, as previously noted (35). Adapting the model to our task, the total cost for performing a lick bout reflects a sum of two terms (**Fig. 3a**). The first term is the energetic cost, reflecting the physical effort in performing actions in quick succession. The form of the energetic cost is assumed to be inversely proportional to the latency between actions (34). The second term is the opportunity cost of forgoing a reward by not performing an action. This cost grows linearly as a function of latency between actions, with a rate for water rewards (*k*_*o*_) and scaling factors to account for accelerated rates for other reinforcement types (*c*_*3nc*_, *c*_*3pc*_, *c*_*10*_). There is an additional factor in the second term for marginal utility, which captures the satiety of the animal after acquiring a certain number of rewards (*R*_*o*_) and the subsequent exponentially diminishing returns (*α*). The goal of the animal is to select the appropriate latency for the self-paced action to minimize the total cost. By calculating the expected latencies, which change across a session due to varying reinforcement type and increasing satiety level, we may then predict the response rates which equate to the rewards earned on a block-by-block basis.

**Figure 3:**
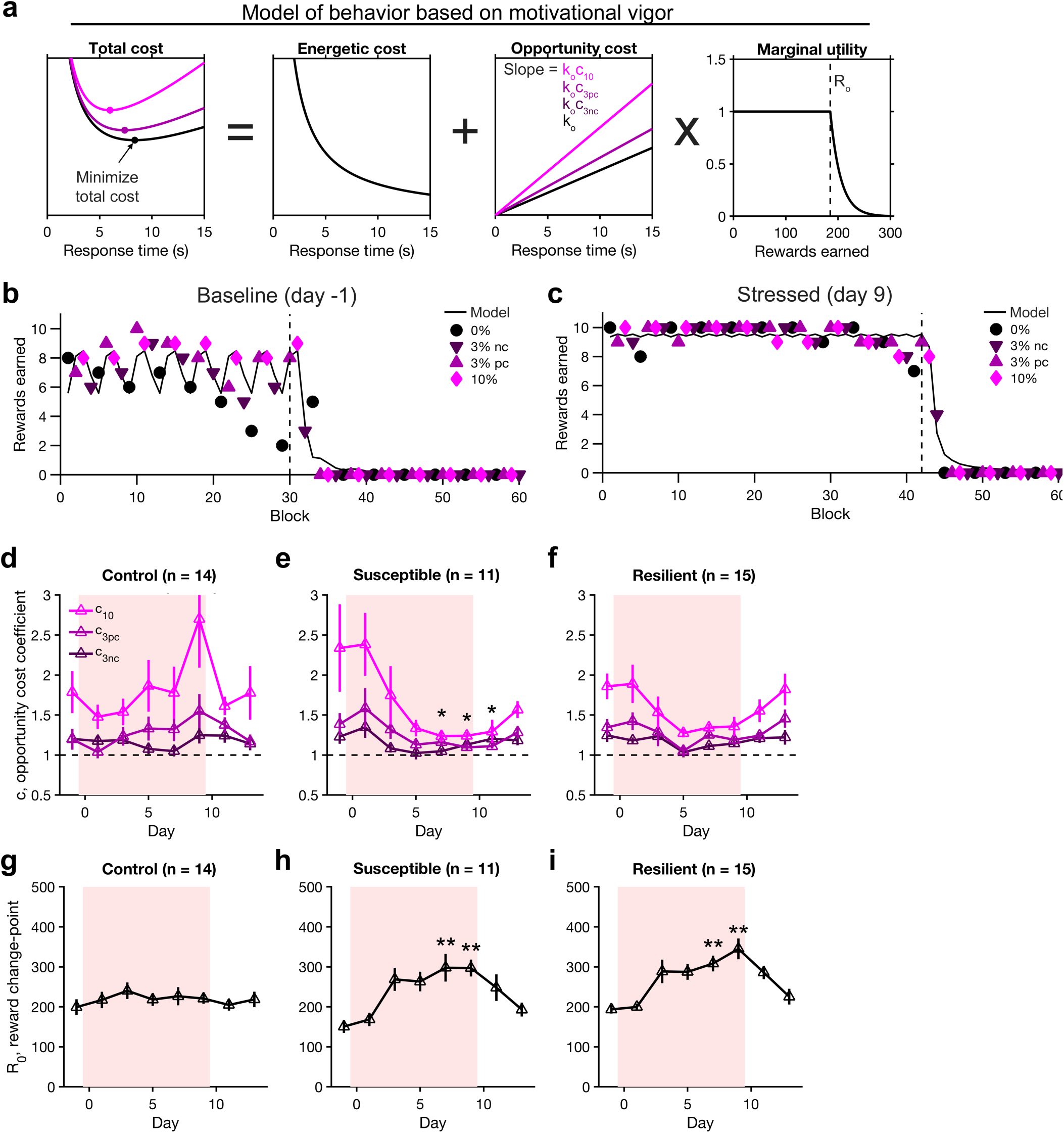
A model of self-paced, instrumental sucrose preference based on motivational vigor. **(a)** Schematic of the model. Animal should select a response time that minimizes the total cost. Total cost is the sum of an energetic cost, incurred if actions have to be performed in quick succession, and an opportunity cost, incurred when potential rewards are delayed due to inaction. The opportunity cost depends on the expected reinforcement type, and a marginal utility term for diminishing returns due to satiation. From the optimal response time, we could estimate the number of completed actions per block. **(b)** Task performance and model fit for an example session on day -1. **(c)** Same as **(b)** on day 9 for the same animal. **(d)** The opportunity cost coefficients determined by fitting model to individual sessions for control animals. Line, mean ± s.e.m. Shading, days with social defeat. Main effect of reinforcement type (F_2,305_ = 14.0, *P* = 2 × 10^−6^), but not for day (F_11,305_ = 1.33, *P* = 0.2) or interaction (F_22,305_ = 0.541, *P* = 1), ANOVA with reinforcement type and day as fixed factors, subject as random factor. **(e)** Same as **(d)** for susceptible animals. Main effect of reinforcement type (F_2,269_ = 8.43, *P* = 3 × 10^−4^) and day (F_10,269_ = 4.82, *P* = 3 × 10^−6^), but not for interaction (F_20,269_ = 0.983, *P* = 0.48), ANOVA with reinforcement type and day as fixed factors, subject as random factor. Comparisons: for the c_10_ parameter, versus day -1, *P* = 1.0 for day 1; *P* = 1.0 for day 3; *P* = 0.06 for day 5; *P* = 0.04 for day 7; *P* = 0.02 for day 9; *P* = 0.04 for day 11; *P* = 0.7 for day 13, from post hoc Tukey-Kramer test. **(f)** Same as **(d)** for resilient animals. Main effect of reinforcement type (F_2,344_ = 13.8, *P* = 2 × 10^−6^) and day (F_11,344_ = 3.39, *P* = 2 × 10^−4^), but not for interaction (F_22,344_ = 0.544, *P* = 1.0), ANOVA with reinforcement type and day as fixed factors, subject as random factor. Comparisons: for the c_10_ parameter, versus day -1, *P* = 1.0 for day 1; *P* = 1.0 for day 3; *P* = 0.1 for day 5; *P* = 0.4 for day 7; *P* = 0.5 for day 9; *P* = 1.0 for day 11; *P* = 1.0 for day 13, from post hoc Tukey-Kramer test. **(g)** The change-point parameter for the marginal utility, determined by fitting model to individual sessions for control animals. Line, mean ± s.e.m. Shading, days with social defeat. No effect of day (F_11,101_ = 0.416, *P* = 1), ANOVA with day as fixed factor, subject as random factor. **(h)** Same as **(g)** for susceptible animals. Main effect of day (F_10,89_ = 3.89, *P* = 3 × 10^−4^), ANOVA with day as fixed factor, subject as random factor. Comparisons: versus day -1, *P* = 1.0 for day 1; *P* = 0.07 for day 3; *P* = 0.07 for day 5; *P* = 0.008 for day 7; *P* = 0.004 for day 9; *P* = 0.2 for day 11; *P* = 1.0 for day 13, from post hoc Tukey-Kramer test. **(i)** Same as **(g)** for resilient animals. Main effect of day (F_11,114_ = 5.00, *P* = 3 × 10^−6^), ANOVA with day as fixed factor, subject as random factor. Comparisons: versus day -1, *P* = 1.0 for day 1; *P* = 0.08 for day 3; *P* = 0.05 for day 5; *P* = 0.0006 for day 7; *P* = 0.0001 for day 9; *P* = 0.1 for day 11; *P* = 1.0 for day 13, from post hoc Tukey-Kramer test. Sample sizes were 14 mice **(d, g)**, 11 mice **(e, h)**, 15 mice **(f, i)**. * *P* < 0.05, ** *P* < 0.01, *** *P* < 0.001.

For each session, we fit the model with 5 free parameters (*k*_*o*_, *c*_*3nc*_, *c*_*3pc*_, *c*_*10*_, *R*_*o*_) to the behavioral data. One parameter (*α*) was not part of the fit, because its value was assumed to be constant across sessions and it was only fitted once using calibration data (see Methods). For pre-stress sessions, the model reproduced the sawtooth oscillations in the block-by-block reward rates and the eventual cessation of licking (**Fig. 3b**). Moreover, the model was flexible enough to also fit sessions involving motivational anhedonia following social defeat stress (**Fig. 3c**).

The model was useful for distilling the session-by-session variations in behavioral performance into a few parameters. More specifically, the motivational component of action was captured by the opportunity cost coefficients (*c*_*3nc*_, *c*_*3pc*_, *c*_*10*_; **Figs. 3d – f**). Focusing on the susceptible animals, the reward sensitivity was significantly modulated by reward type and day (*P* = 3 × 10^−4^, main effect of reward type; *P* = 3 × 10^−6^, main effect of reward type; ANOVA with block type and day as fixed factors and subject as random factor; **Fig. 3e**). Notably, the motivational deficit did not emerge immediately following the first few stress sessions, but rather became detectable only starting on day 7. The delayed emergence indicates that the motivational anhedonia is a consequence of the cumulative impact of the chronic stressor.

Stress also has a marked influence on the reward change-point for the marginal utility (*R*_*o*_). Control animals attained a stable number of rewards until satiety across behavioral sessions (**Fig. 3g**). However, susceptible and resilient animals had a significantly higher threshold for marginal utility, such that they completed more actions until satiety (**Figs. 3h – i**). The effect was limited to the days with stress exposures, as it quickly fell back to baseline after the last defeat session. Increased reward change-point means increased intake for stressed animals during the task. This agrees with some studies showing that animals subjected to social stress raise their food or fluid intake (36, 37), although the interaction with metabolism is complex (38) and stress-induced changes in intake is not a consistent finding in the literature (14).

### Stress effects on appetitive anhedonia were transient

We next analyzed how repeated social defeat affects consummatory licks, which pertains to the appetitive component of action. In the pre-stress stage, the dependence of behavior on reward can be visualized by plotting single-trial lick raster and trial-averaged lick density for 0%, 3%nc, 3%pc, and 10% blocks. As seen in **Figure 4a**, although there were trial-by-trial variations, there is a robust positive dependence on the reward value. By contrast, the same animal after multiple defeat sessions had qualitatively similar lick timing, but did not show any reward dependence (**Fig. 4b**). Across all control animals, consummatory licking was dependent on the reward type (*P* = 0.002, main effect of reward type; ANOVA with block type and stage as fixed factors and subject as a random factor; **Fig. 4c**), as would be expected. For susceptible animals, the number of consummatory actions was dependent on both reward type and stress stage (*P* = 0.04, main effect of reward type; *P* = 0.04, main effect of stage; ANOVA with block type and stage as fixed factors and subject as a random factor; **Fig. 4d**). Notably, reward sensitivity for consummatory actions was diminished for the days of stress exposures, but quickly rebounded to baseline levels following the last day of stress. The impact of stress on consummatory actions was qualitatively similar, though weaker, for resilient animals (**Fig. 4e**). Overall, these results indicate that this social defeat procedure induced only transient appetitive anhedonia, which differs from the protracted motivational anhedonia in susceptible animals.

**Figure 4:**
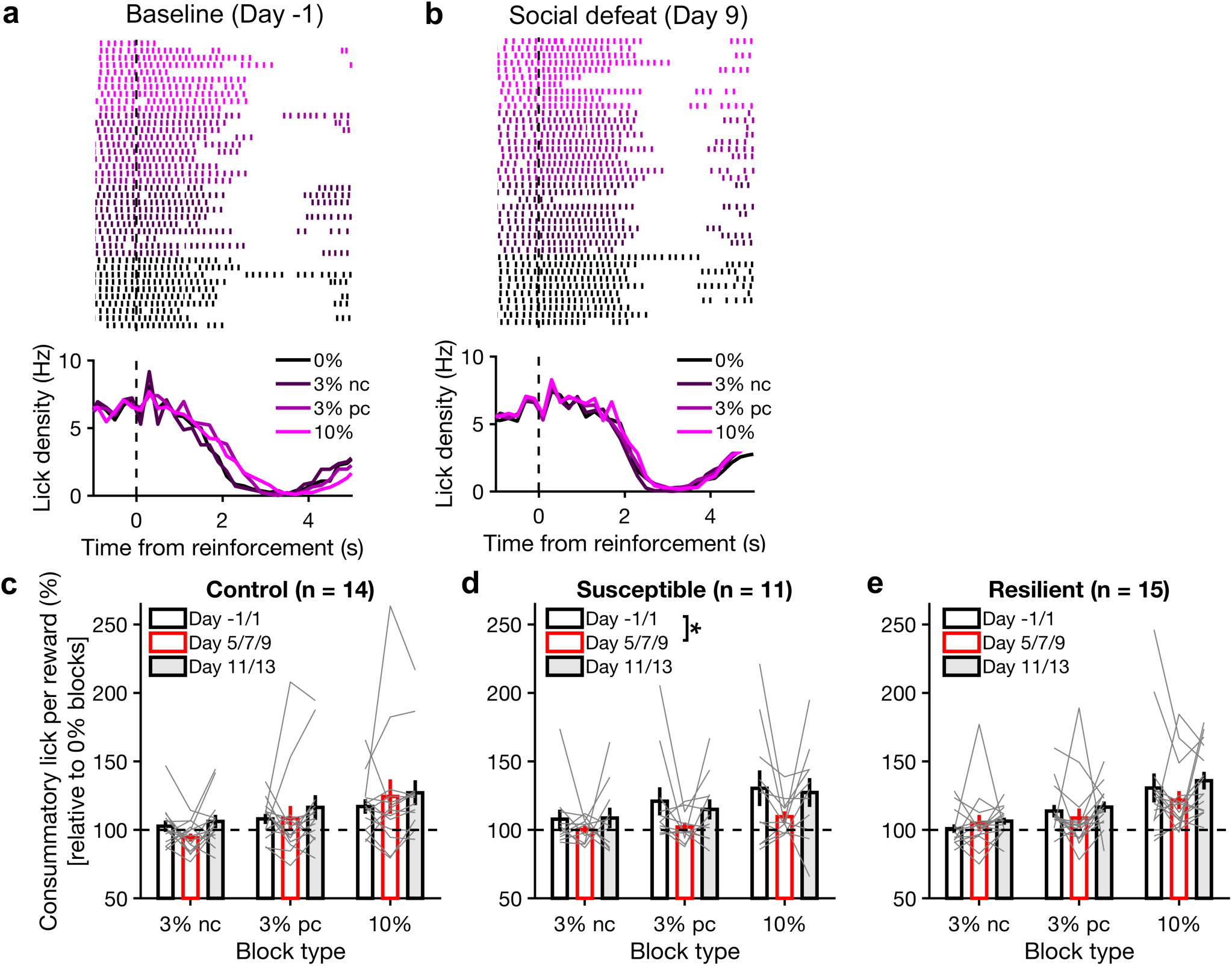
Chronic social defeat had only transient effect on consummatory actions. **(a)** Top, lick raster, chosen randomly from 10 trials of each reinforcement type from a session on day -1. Bottom, mean lick density relative to the time of reinforcement, plotted separately for each reinforcement type, for the same session. **(b)** Same as **(a)** on day 9 for the same animal. **(c)** Consummatory licks per reward for each reinforcement type, normalized by the number in 0% blocks, for control animals. For each animal, the value was an average across days according to three stages: “pre” for day -1 and 1, “stress” for day 5, 7, and 9, and “post” for day 11 and 13. Gray line, individual animal; bar, mean ± s.e.m. Main effect of reinforcement type (F_2,117_ = 6.49, *P* = 0.002), but not for stage (F_2,117_ = 0.99, *P* = 0.4) or interaction (F_4,117_ = 0.28, *P* = 0.9), ANOVA with reinforcement type and stage as fixed factors, subject as random factor. **(d)** Same as **(c)** for susceptible animals. Main effect of reinforcement type (F_2,90_ = 3.35, *P* = 0.04) and stage (F_2,90_ = 3.30, *P* = 0.04), but not interaction (F_4,90_ = 0.22, *P* = 0.9), with comparisons: pre vs. stress, *P* = 0.048; pre vs. post *P* = 0.9; stress vs. post, *P* = 0.2, ANOVA with reinforcement type and stage as fixed factors, subject as random factor, and post hoc Tukey-Kramer test. **(e)** Same as **(c)** for resilient animals. Main effect of reinforcement type (F_2,126_ = 13.3, *P* = 5 × 10^−6^), but not stage (F_2,126_ = 1.27, *P* = 0.3) or interaction (F_4,126_ = 0.39, *P* = 0.8), ANOVA with reinforcement type and stage as fixed factors, subject as random factor. Sample sizes were 14 mice **(c, d)**, 11 mice **(e, f)**, and 15 mice **(g, h)**. * *P* < 0.05.

### Prefrontal Cg1/M2 region as a likely substrate for mediating the effects of repeated stress on self-paced actions

Where should one look for the neural substrate underlying the stress-induced deficit in reward-directed actions? Although motivated behavior is known to involve a broad frontal-basal ganglia network, there is growing evidence that the cingulate (Cg1) and medial secondary motor (M2) regions, which constitute the most dorsal aspect of the rodent medial prefrontal cortex (39), are particularly important for the production of actions that are self-initiated and reward-guided (40-42). Specifically, the timing of self-initiated actions depends on an intact Cg1/M2 (43) and is tracked closely by the activity of individual neurons in the region (44). The function of Cg1/M2 is restricted to goal-directed actions, because when the region is lesioned, animals perform fewer actions in a ratio schedule, but habitual responding in an interval schedule remains intact (45). Furthermore, Cg1/M2 is a target of stress, as identified by brain-wide mapping of neuronal activation in a rodent model of depression (46). Cg1/M2 is also engaged following systemic administration of the fast-acting antidepressant ketamine (47), which enhances synaptic calcium signaling and promote dendritic remodeling in this brain region (48, 49). Altogether, because Cg1/M2 is essential for the type of self-paced and fixed-ratio responses employed in our task, and because Cg1/M2 is targeted by stress and fast-acting antidepressant, we investigated how neural activity dynamics in this subregion of the medial prefrontal cortex responds to repeated stress.

### Imaging the time course of neural activity changes at cellular resolution

During a task, if we were to observe altered neural activity in stressed animals, the results could arise from the direct effect of stress on neurons or the indirect consequence of stress modifying the animal’s responding during behavior. Therefore, to determine the direct effect of stress on neural activity, we measured spontaneous neural activity in the absence of behavior. Neuronal firing leads to somatic calcium transients in pyramidal neurons (50), which could be visualized using a fluorescent calcium indicator. To characterize activity changes across days, we used a two-photon microscope to track and record from the same set of cells. One challenge for longitudinal calcium imaging is that the fluorescence signal is activity-dependent, making it difficult to identify the same cells unequivocally every session. To overcome this challenge, we used a recently developed bicistronic vector to express the genetically encoded calcium indicator GCaMP6s along with a static fluorophore mRuby2 (51). Targeting Cg1 and medial M2 (AP=1.5 mm, ML=0.4 mm), we injected AAV-hSyn1-Flex-mRuby2-GSG-P2A-GCaMP6s-WPRE-pA in CaMKIIa-Cre mice for selective expression in pyramidal neurons (**Fig. 5a**). A glass window was chronically implanted for long-term optical imaging. Histology and *in vivo* imaging confirmed overlapping expression of GCaMP6s and mRuby2 in Cg1/M2 (**Fig. 5b – c**). Using static mRuby2 fluorescence to identify cell assemblies, we could track individual neurons in layer 2/3 of Cg1/M2 with a high degree of certainty for up to 7 sessions spanning 13 days (**Figs. 5d – e**). We used CaMKIIa::GCaMP6s-mRuby2 animals for the majority of the longitudinal imaging experiments. A smaller subset of data came from earlier experiments with C57BL/6J animals injected with AAV-CaMKII-GCaMP6f. Although yield from longitudinal tracking in the earlier experiments was lower, there was no obvious difference in the functional characterization, therefore we combined the data and present the collective results.

**Figure 5:**
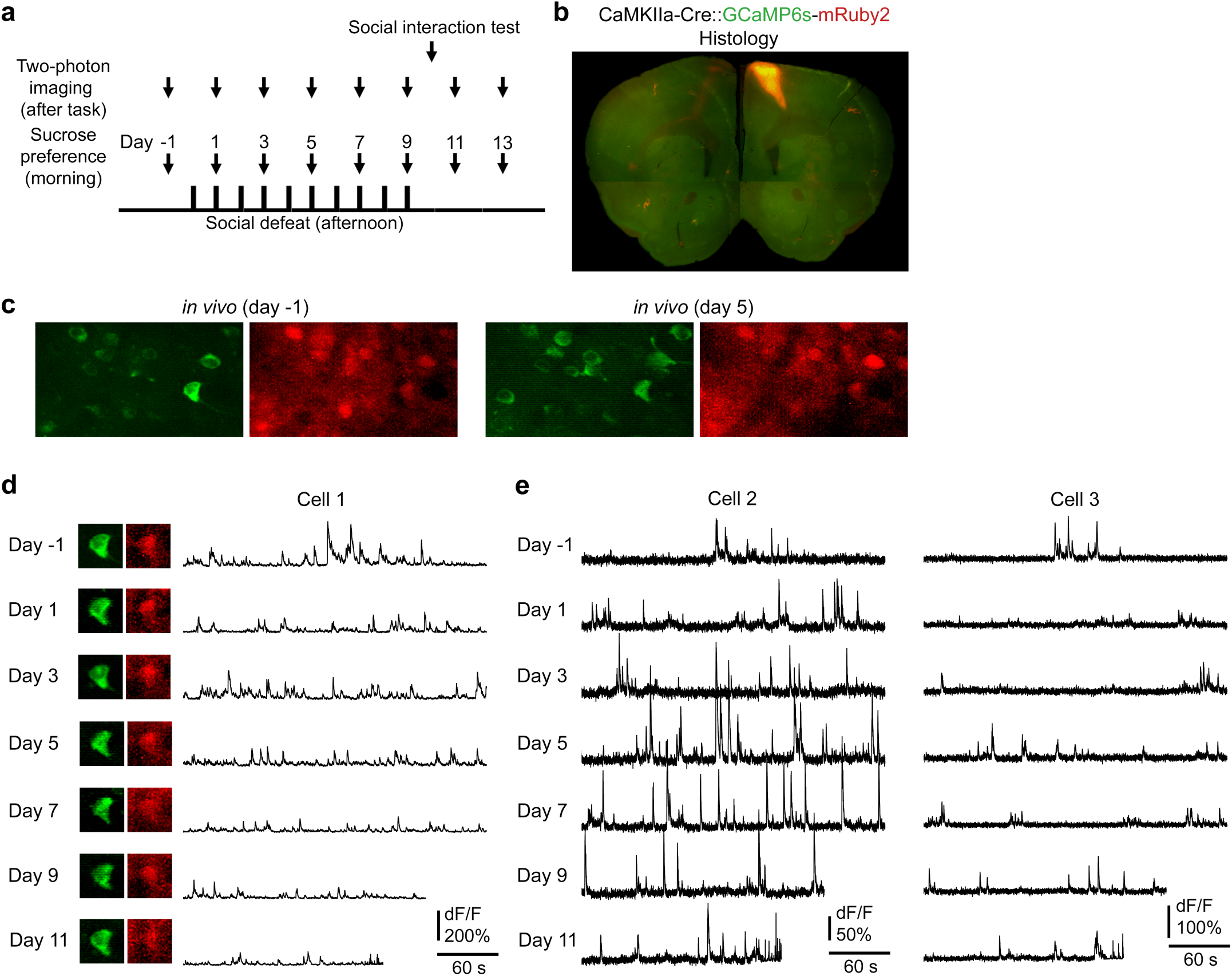
Longitudinal imaging of spontaneous activity of layer 2/3 pyramidal neurons in Cg1/M2. **(a)** Timeline of the experiments. **(b)** A fixed coronal section showing the extent of virally mediated expression of GCaMP6s-mRuby2 in Cg1/M2. **(c)** *In vivo* two-photon images from an awake, head-fixed mouse for GCaMP6s (green) and mRuby2 (red) in pyramidal neurons in Cg1/M2. The images were taken 6 days apart. **(d)** Spontaneous fluorescence transients for an example cell from (c), recorded 2 days apart across 7 sessions spanning 13 days. The insets show a still frame of the *in vivo* GCaMP6s and mRuby2 fluorescence on the corresponding days. **(e)** Similar to (d) for two other cells.

### Social defeat stress initially elevates the activity of prefrontal pyramidal neurons

We devised an experimental plan to include two parts. For the first part, we characterized the short-term effects of a single defeat session on neural activity with stressed (n = 7) and control (n = 4) animals (**Fig. 6a**). For the second part, focusing our effort to determine the effects of repeated stress, we tracked the long-term effects of multiple defeat sessions in stressed animals. The expectation was that stressed animals would separate into susceptible and resilient subtypes, allowing for a comparison of the neural activity changes between phenotypes. Indeed, the social interaction test conducted after the last day of social defeat indicated that, out of the 7 stressed mice in the long-term imaging experiment, 4 animals were classified as susceptible and 3 animals were classified as resilient (**Fig. 6b**). The result indicates that the chronic social defeat stress had the expected behavioral impact for imaging mice.

**Figure 6:**
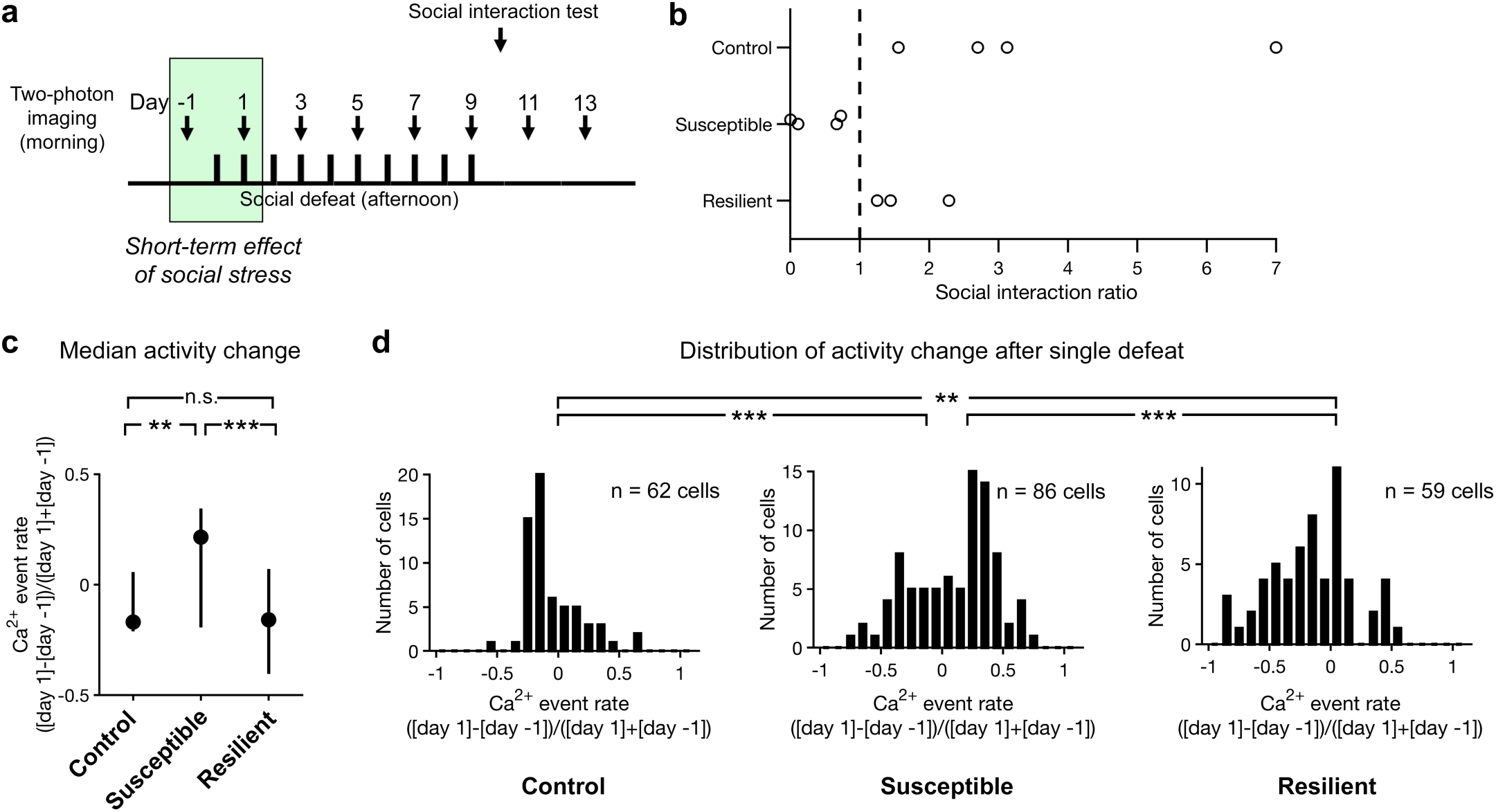
Social defeat rapidly alters the prefrontal cortical activity. **(a)** The timeline for experiments to determine the impact of social defeat stress on prefrontal cortical activity. The shaded green area represents the time window used for analysis of the short-term effects. **(b)** Based on the social interaction ratio, mice subjected to social defeat were divided into resilient and susceptible individuals. Controls were handled but did not experience social defeat. Each circle represents an animal. Note that the imaging animals were a new cohort and not part of the behavior-only study. **(c)** Median change in neural activity from day -1 to 1 for control, susceptible, and resilient animals. The activity change was a normalized difference in the rate of events inferred from fluorescence transients, calculated separately for each cell. Filled circles, median; bar, 25^th^ and 75^th^ percentiles. Control versus susceptible: *P* = 0.002, control versus resilient: *P* = 0.3, susceptible versus resilient: *P* = 0.0001, Wilcoxon rank-sum test. **(d)** Histogram of the neural activity change from day -1 to 1 for control, susceptible, and resilient animals. Control versus susceptible: *P* = 2.1 × 10^−5^, control versus resilient: *P* = 0.005, susceptible versus resilient: *P* = 6.1 × 10^−6^, Kolmogorov-Smirnov test. Sample sizes for **(c)** and **(d)** were 62 cells from 4 control animals, 86 cells from 4 susceptible animals, and 59 cells from 3 resilient animals. ** *P* < 0.01, *** *P* < 0.001.

To determine the short-term effect of social stress on Cg1/M2, we compared the spontaneously occurring calcium transients in layer 2/3 pyramidal neurons in awake mice, one day before versus one day after a single defeat. To infer activity rates from somatic fluorescence signals, we used a peeling algorithm based on template matching that has been extensively validated in prior studies (52, 53) (see Methods). When we examined the overall activity change across all cells, susceptible animals had a significant elevation of spontaneous activity following a single defeat, relative to control animals (control versus susceptible: *P* = 0.002, Wilcoxon rank-sum test; **Fig. 6c**). Intriguingly, resilient and control animals were indistinguishable by this measure of median activity change (control versus resilient: *P* = 0.3, Wilcoxon rank-sum test). However, when we plotted the distribution of activity changes, a more nuanced picture arose. The distribution of the activity change was more dispersed for susceptible and resilient animals (interquartile range: 0.27 for controls, 0.54 for susceptible, 0.47 for resilient; **Fig. 6d**). Specifically, for resilient animals, this means that even though the median did not change, there were more cells with greater decrease or increase in spontaneous activity following stress relative to controls (control versus resilient: *P* = 0.005, Kolmogorov-Smirnov test). Collectively, these results demonstrate that a single stress episode already had a detectable impact on activity of layer 2/3 pyramidal neurons in Cg1/M2, with particularly pronounced hyperactivity for susceptible animals. Moreover, there is a resilience-specific adaptation that maintained the overall activity to be comparable to control animals, but was revealed at the level of ensemble activity.

### Social defeat stress exerts heterogeneous long-term effects on prefrontal cortical activity

We continued to image the stressed animals and, for a majority of the cells, we were able to track their activity throughout and beyond the social defeat manipulation (**Fig. 7a**). Using heatmaps, we could visualize and compare how the spontaneous activity levels of layer 2/3 pyramidal neurons in Cg1/M2 changed relative to pre-stress baseline (**Fig. 7b**). For susceptible animals, many neurons had an initial elevation in spontaneous activity after the first defeat session. However, with more stress exposures, the activity changes became more heterogeneous. Some Cg1/M2 neurons had sustained elevated activity, whereas many others had a marked decrease to below baseline following chronic stress. By contrast, for resilient animals, the effects of repeated social defeats on prefrontal cortical activity was more similar across neurons, amounting mostly to a reduction in spontaneous activity.

**Figure 7:**
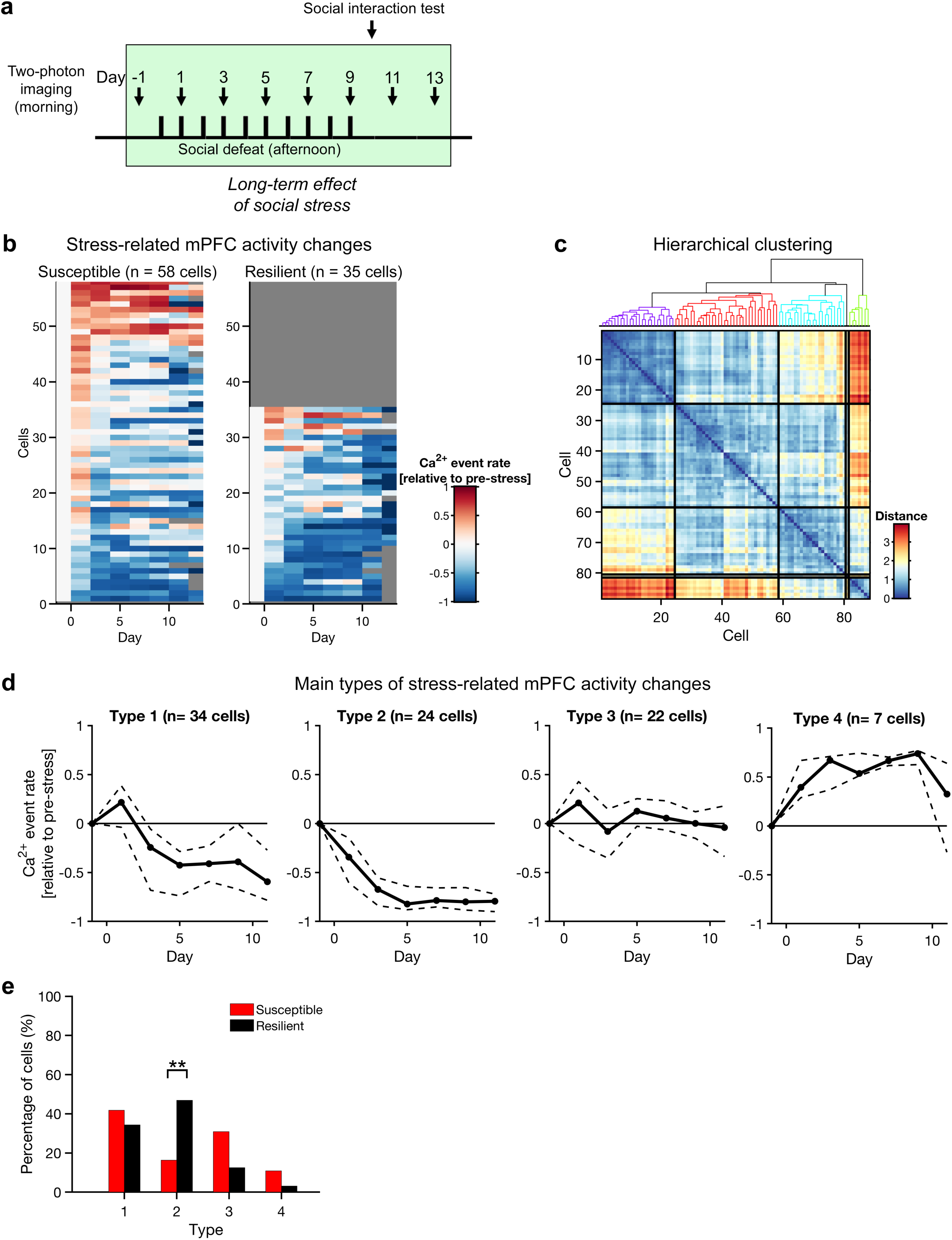
Social defeat induces heterogeneous long-term activity changes in the prefrontal cortex. **(a)** The timeline for experiments to determine the impact of social defeat stress on prefrontal cortical activity. The shaded green area represents the time window used for analysis of the long-term effects. **(b)** Heatmaps showing the change in spontaneous activity relative to the pre-stress baseline. Each row is a cell. The left and right heatmaps contain the 58 cells from 4 susceptible animals and 35 cells from 3 resilient animals respectively. Red indicates an increase in spontaneous activity relative to pre-stress day -1. Blue indicates a decrease. **(c)** Results of applying hierarchical clustering with Euclidean distance as the distance metric on the entire data set of 93 cells. **(d)** Cells were classified as Type 1 – 4 based on the cluster membership identified by the hierarchical procedure. For each Type, the change in spontaneous activity was averaged across cells. Solid line, median. Dotted lines, 20^th^ and 80^th^ percentiles. **(e)** The proportion of cells in susceptible or resilient animal that was classified for each Type. Comparing the proportions between susceptible and resilient animals, the difference was significant for Type 2 (*P* = 0.002, chi-square test), and did not reach significance for Type 1 (*P* = 0.49), Type 3 (*P* = 0.05), and 4 (*P* = 0.20).

To delineate the time courses of stress-induced activity changes more systematically, we applied hierarchical clustering to the entire sample of 93 neurons, including 58 cells from susceptible animals and 35 cells from resilient animals. For all possible pair of cells, the similarity between their day-by-day activity changes was determined by calculating differences in Euclidean distance. The clustering procedure then agglomerated similar cells into four major clusters (see Methods). The quality of the clusters, in terms of their distance separation, could be assessed visually by dendrogram and pairwise similarity matrix (**Fig. 7c**). We denoted pyramidal neurons belonging to the four major clusters as Type 1, 2, 3, and 4. These functional cell types had distinct time courses (**Fig. 7d**), and could be found at different prevalence in susceptible and resilient animals (**Fig. 7e**). Type 1 cells included neurons that had an initial elevation of spontaneous activity, followed by decreases for the next two sessions down to a sustained level below the baseline. This was the most frequently observed type of time course and cells with this profile could be found in both susceptible and resilient animals (*P* = 0.49, chi-square test). Type 2 cells had a monotonic decline in spontaneous activity as a response to repeated stress. The decrease continued across the initial stress sessions before stabilizing to a steady below-baseline level. Notably, Type 2 cells were significantly more common in resilient animals (*P* = 0.002, chi-square test). By contrast, Type 3 and Type 4 cells were more abundant in the susceptible animals, although the comparisons did not reach statistical significance likely because there were few cells with these time courses (Type 3: *P* = 0.05, Type 4: *P* = 0.20, chi-square test). Type 3 and Type 4 cells had a time course that was either steady near the baseline or had prolonged heighted spontaneous activity. In particular, Type 4 cells might be the most related to the stress-induced motivational anhedonia, as they were predominantly found in susceptible animals (6/7 cells) and mirrored the progression of the behavioral deficit. These results show the heterogeneous impact of social defeat on prefrontal cortical activity. Moreover, for the majority of neurons, the activity modification evolves across repeated stress sessions, highlighting the cumulative nature of stress impact at the level of neural activity.

## Discussion

In this study, we measured the longitudinal effects of repeated social defeat on reward-directed actions and determined the concomitant profiles of activity changes for layer 2/3 pyramidal neurons in Cg1/M2. There are two main findings. First, behavioral and neural deficits in response to social defeat build over multiple stress episodes. This suggests that the response to repeated stress is not a unitary behavioral condition, but rather represents a continuum where accumulating exposure progressively deteriorates reward processing. The progression was observed in behavior as well as in activity of individual neurons. Second, resilience is associated with neural activity adaptations that are not only distinct from susceptible mice, but also dissimilar from control animals. This finding raises the possibility that resilience involves the capacity to reorganize prefrontal cortical activity adaptively to counteract the negative impact of social stress.

We have attributed specific performance metrics – the number of earned rewards and consummatory licks – to the motivational and appetitive components of the animal’s behavior. The validity of this analysis is supported by a large number of prior studies. For the motivational component, a fixed or random ratio schedule forces an animal to perform a prespecified number of responses to obtain a reward. This type of schedule biases subjects towards performing goal-directed actions over habitual responding, as has been demonstrated by classic experiments involving reinforcer devaluation (23). For the appetitive component, it has long been appreciated that the number of licks in a bout has a positive, monotonic relationship with the concentration of palatable fluids (29). As a result, along with supporting evidence involving substituting with unpalatable fluids, pairing palatable taste with lithium chloride, and administering benzodiazepine, the lick cluster size has shown to reflect an animal’s appetitive reaction (30).

One limitation of the current behavioral setup is that both the instrumental and consummatory actions in this task are mediated by licking. For future studies, it may be possible to replace the instrumental responses for reward with another apparatus, for example a lever. In pilot experiments, we have tested using a front-paw press on a snap-action lever switch as the instrumental response. However, we find that although the mouse can press the lever, their movements vary and they do not press many times on a fixed-ratio schedule, perhaps because the action is more physically demanding for a head-fixed animal. Instead, to isolate consummatory licks, we have introduced a 5-second timeout period following each reinforcement. Mice learn to cease licking during this period, and therefore the number of consummatory licks is not contaminated by the subsequent instrumental actions that belong to a different lick bout (**Figs. 4a – b**).

Using the self-paced, instrumental sucrose preference task, we show that chronic social defeat stress leads to motivational anhedonia. This reduced reward preference was not due to reduced intake of the sucrose solutions, but rather it was due to increased and indiscriminate responding for all solutions (**Fig. 3e, h**). This is broadly consistent with previous studies demonstrating a stress-induced shift from goal-directed actions to habitual responding in humans and rodents (7, 8). A few other studies have charted the time course of stress-induced effects on motivation, such as with intra-cranial self-stimulation in rats (54, 55). In one study, the diminished reward sensitivity rises sharply following stress onset for all animals, but only is only protracted for susceptible subjects (54). By contrast, here we report a delayed emergence of the motivational deficit for both susceptible and resilient animals. The discrepancy may be due to the reduced sensory interaction time in our protocol, which probably decreases the intensity of the stressor, and allows for a clearer view of the cumulative progression.

On average across all cells, we showed an initial elevation and then subsequent reduction of spontaneous activity in prefrontal pyramidal neurons in susceptible animals. These results match well with what we know in terms of stress-induced glutamatergic dysfunction in the prefrontal cortex (56). In particular, acute restraint stress leads to an immediate increase in extracellular glutamate recorded in the medial prefrontal cortex, which peaks 20 min after the termination of stress but continues to be elevated up to the latest data point measured at 80 min post-stress (57) This initial hyperexcitation may relate to the increased surface levels of glutamatergic receptors, as well as potentiated synaptic responses following acute exposure to stress (58). Studies of chronic stress exposure have generally showed the opposite trend of prefrontal cortical hypoactivity. Mice subjected to 10 days of social defeat have reduced glutamate levels in the prefrontal cortex (31). A number of cellular mechanisms may underlie the reduced glutamatergic activity, including the loss of AMPA and NMDA receptors (17), structural atrophy such as dendritic retraction and synapse loss (15, 16), and maladaptive synaptic inhibition (59).

The novelty of this study is that we have measured the progressive change in cellular activity as the animal experiences successive stress episodes and develops a behavioral response to that stress. There is considerable heterogeneity in how individual prefrontal cortical neurons respond as a function of stress. Previous studies have hinted at heterogeneous neural responses on a short-term time scale. For example, it has been reported that prefrontal cortical neurons can have either transient or protracted firing responses to restraint (60). Moreover, stress-related excitatory synaptic modifications may be selective to a subset of prefrontal cortical neurons (18). In the current study, we used hierarchical clustering to identify several major profiles of stress-induced activity changes in prefrontal cortical neurons. The most numerous neuronal subtype (Type 1) display bidirectional activity changes involving initial hyperactivity followed by sustained hypoactivity. This time course might be expected based on the glutamatergic dysfunction associated with acute and chronic stress. Interestingly, both susceptible and resilient animals had a large number of Type 1 neurons, suggesting that this functional cell type may not be the distinguishing feature of susceptibility versus resilience. Instead, Type 2, 3, and 4 neurons have a monotonic progression of activity changes over time, and their proportions differ depending on the animal’s reactivity to social stress. The implication is that a minor fraction of prefrontal cortical neurons may be particularly relevant for mediating the stress-induced behavioral deficits.

In addition to the long-term differences in time courses, we demonstrated distinct features in neural activity changes following a single defeat between susceptible and resilient animals. Intriguingly, resilient animals were different from both susceptible and control animals. At the ensemble level, resilient animals were characterized by a wide range of activity changes, yielding a median activity change that is comparable to control animals but significantly lower than susceptible mice. These observations suggest that resilience is associated with adaptations of prefrontal cortical activity that brings the overall equilibrium to a normal level. By contrast, susceptibility to social stress may represent a failure to reorganize neural activity appropriately to counteract the stress effects, leading to a hyperactive state. More experiments will be needed to relate these findings to other molecular and network mechanisms that have been suggested to confer vulnerability and resilience to social stress (61, 62).

To sum, the results presented here demonstrate a progressive development of behavioral and neural deficits in response to an ethologically relevant social stress. The data provide direct support for the allostatic load hypothesis, which posits that maladaptations arise from the cumulative burden of repeated stress (2). Looking ahead, a number of studies have shown that depressive-like phenotypes in rodents may be reversed by inducing synaptic plasticity in the medial prefrontal cortex. This may be achieved by controlling firing rates using optogenetics (11, 63) or by the administration of fast-acting antidepressants such as ketamine (19, 48, 64). Except in a few cases (65, 66), to date most studies have manipulated prefrontal pyramidal neurons indiscriminately. By showing that individual cells can have distinct profiles of activity adaptation in response to social stress, our findings suggest that therapeutic interventions that target functional subtypes of pyramidal neurons may approximate more closely the physiological process of resilience and therefore be effective at reversing behavioral deficits.

## Methods

### Animals

We used male, adult (2 – 6-month-old) mice. Behavioral experiments were performed using wild type C57BL/6J mice (#000664, Jackson Laboratory). Imaging experiments were performed using C57BL/6J mice or heterozygous CaMKIIa-Cre mice (#005359, Jackson Laboratory). For chronic social defeat stress, aggressive residents were CD-1 male retired breeders (Crl:CD1(ICR), #022, Charles River).

### Surgery

Anesthesia was induced by isoflurane at 2% in O_2_ and maintained for the duration of the surgery at 1 – 1.5% in O_2_. Anesthesia level was monitored by visual observation of the breathing rhythm and by the lack of response to paw pinches. The animal was placed on a water-circulating pad (TP-700, Gaymar Stryker) set at 38 °C to prevent hypothermia. The head was positioned in a stereotaxic frame (Kopf Instruments). The animal received preoperative care including carprofen (5 mg/kg, s.c., Butler Animal Health) and dexamethasone (3 mg/kg, s.c., Henry Schein Animal Health). A midline incision was made to expose the skull. A stainless-steel head plate (eMachineshop.com), necessary for head fixation, was then affixed to the skull with cyanoacrylate glue (Loctite 454, Henkel) and dental cement (C&B Metabond, Parkell). After surgery, the mouse was given time in isolation to recover in a recovery cage for an hour and then return to the home cage. After each surgery, the mouse received post-operative care including injections of carprofen (5 mg/kg, s.c.) and dexamethasone (3 mg/kg, s.c.). One dose was given at the end of the surgery and then one dose each day for the 3 subsequent days. The mouse recovered for at least 1 week in home cage before experiments began.

For experiments involving calcium imaging, we performed additional steps after the midline incision and before the head plate implantation. We targeted the Cg1 and M2 regions of the medial prefrontal cortex (AP = +1.5 mm and ML = +0.4 mm relative to bregma). A 3-mm diameter craniotomy (centered at AP = +1.5 mm and ML = +1.5 mm relative to bregma) was made over the right hemisphere using a dental drill. The dura was kept moist with sterile saline solution (0.9% Sodium Chloride, preservative free, Hospira). To express GCaMP6s in pyramidal neurons, we injected pAAV(AAV1)-hSyn1-Flex-mRuby2-GSG-P2A-GCaMP6s-WPRE-pA (Addgene) in CaMKII-Cre mice. This bicistronic construct was created to include a static fluorophore for the purpose of longitudinal calcium imaging (51). In a subset of experiments, to express GCaMP6f in pyramidal neurons, we injected pAAV(AAV1)-CaMKII-GCaMP6f-WPRE-SV40 (Penn Vector Core or Addgene) in C57BL/6J mice. For each animal, we injected 184 nL at a depth of 0.4 mm from dura using a glass pipette mounted on a microinjector (Nanoject II, Drummond). A custom glass plug was fabricated by gluing together four layers of #1 thickness 3-mm diameter round coverslips (64-0720 CS-3R, Warner Instruments) topped with one layer of #1 thickness 4-mm diameter round coverslip (64-0724 CS-4R, Warner Instruments), using an optical adhesive (61, Norland Optical Adhesive) cured by exposure to LED 365 nm light (Loctite EQ CL32 LED Spot 365 nm, Henkel). Following virus injection, the glass plug was placed to cover the craniotomy, such that the bottommost 3-mm coverslip was in contact with the dura. The glass plug was secured to the skull by applying a ring of cyanoacrylate glue (Loctite 454, Henkel) on the border of the top-layer 4-mm coverslip. The mouse recovered for at least 3 weeks in home cage to allow for viral-mediated expression of the transgene.

### Self-paced, instrumental sucrose preference task

The task design was inspired by prior studies involving ratio schedules (23), incentive contrast (24, 25), and head fixation for awake, mobile mice (26). For the behavioral setup, the mouse was head-fixed but could move its body freely on a treadmill, either a skewered Styrofoam ball or a 7”-diameter plexiglass running disk, to reduce the stressful effect of restraint. Head fixation was achieved by screwing the implanted metal head plate on an angled metal bar itself fixed on the table. Treadmill could freely rotate forward or backward in response to the mouse’s movements. The mouse was positioned in front of a lick spout made by soldering together 3 blunted 20-gauge hypodermic needles (BD PrecisionGlide, 305175). Each lick spout was connected to its own solenoid valve (MB202-VA30-L204, McIntosh Controls), allowing independently delivery of one of three solutions: 0% sucrose (i.e., tap water), 3% sucrose (w/v), and 10% sucrose. The volume delivered by each needle was calibrated to be 4 µL. A tongue lick contact with the lick spout was detected via the closing of an electrical circuit. Lick detection and valve opening were controlled by Presentation software (Neurobehavioral Systems), through two independent data acquisition boards (USB-201, Measurement Computing). To motivate the animal to perform the task, the mouse was fluid restricted. On alternate days, animals would either obtain its entire fluid intake from the task, or be given 15 minutes of ad libitum access to water. To shape the mouse to perform the task, after an initial period of 24 hours of water restriction, the mouse was placed in the behavioral setup to learn the association between licking the spout and fluid delivery. During this initial shaping phase, each lick triggered the delivery of 4 µL of fluid. The fluid type was randomly interleaved, such that 0% sucrose, 3% sucrose, and 10% sucrose solutions were selected with equal probabilities. Typically, the mouse would learn to lick within the first session, as attested by the high number of rewards collected. Nevertheless, to ensure that the mice correctly learned to lick the spout and habituate to the fluid type, the mouse would be trained on two consecutive days for 1 hour each. After this initial shaping phase, the mouse was tested on the self-paced, instrumental sucrose preference task. During the testing phase, the mouse was tested every other day. On test day, the mouse received fluids only during the task. On non-test day, the mouse had 15 minutes of *ad libitum* water access in the home cage. The task had a tandem fixed-ratio-10, fixed-interval-5 schedule of reinforcement (FR10-FI5). Namely, the mice must make 10 licks onto the spouts (FR-10) to receive 4 µL of fluid reinforcement. Following the fluid delivery, there was a 5-second timeout period (FI5) during which licks do not count towards next FR10. Following this 5-second period, the mouse would have to make another 10 licks to trigger the delivery of another fluid reinforcement, and so forth. The reinforcement type changed over the course of a session. At the beginning of a session, after completion of the first FR10, the mouse would receive 4 µL of 0% sucrose, and then has 1 minute to perform more licks to acquire more of the same 0% sucrose reinforcements. After the end of the 1-minute block, the completion of the next FR10 would lead to 4 µL of 3% sucrose, and the mouse would have 1 minute to acquire more of that type of reinforcement, and so forth. The reinforcement type cycles from 0%, 3%, 10%, to 3%, and then again from 0%, 3%, 10%, to 3%, and so forth. The design enabled us to compare behavior across 1-minute blocks with absolute difference in sucrose concentration (e.g., 0% versus 10%), or with relative incentive contrast (e.g., 3% after 0% for positive contrast versus 3% after 10% for negative contrast). To test how the block design influences licking behavior, a separate cohort of animals was tested on a task variant, still employing FR10-FI5, but with no blocks and reinforcements were chosen randomly after each FR10 with each reinforcement type having equal probabilities. To test effects related to the caloric content of sucrose, a separate cohort of animals was tested on a test variant in which the reinforcement types were sucralose, a non-caloric artificial sweetener, at concentrations of 0, 0.23, and 1 mM.

### Repeated social defeat

We followed published procedures to induce social defeat stress (33). Briefly, the protocol uses a partitioned cage, in which the cage space is separated into two halves by a divider made of metal mesh sheet. CD1 retired breeder mice were screened for aggressive behavior and housed in the partitioned cage, allowing the CD1 target to be the resident and the mouse to be stressed to be the intruder. Each day, the intruder was placed into the partitioned half that contains the resident for 10 min to induce bouts of physical aggression. Subsequently, the intruder was placed for another 10 min in the other half of the partitioned cage, allowing for sensory interaction but no physical interaction. Afterwards, the intruder was returned to its home cage with its littermates. This procedure was repeated every day for 10 consecutive days. Each day, the intruder encountered a new, unfamiliar resident. Social defeats was always performed in the afternoon, whereas behavioral tests were always done in the morning, in order to avoid the effects of immediate stress responses in behavioral measurements.

### Social interaction test

We used an open-field chamber with a white floor and plexiglass walls (42 cm (w) × 42 cm (d) × 38 cm (h)). A wire-mesh enclosure, placed against one wall, was used to contain a target CD-1 mouse (8 cm (w) × 6 cm (d) × 38 cm (h)). An overhead camera (SciCam, Scientifica) and infrared illumination (IR30 WideAngle IR Illuminator, CMVision) were used to capture videos of the animals in the dark. For the test, the intruder mouse to be tested was placed in the chamber in the absence of the CD-1 target for 150 s. The intruder mouse was then replaced in the chamber when the CD-1target was present for an additional 150 s. MATLAB scripts were used to extract the animal’s position from the video automatically to determine its trajectory. The social interaction ratio was defined as the time spent in the interaction zone when the CD-1 target was present divided by the time spent in the interaction zone when the CD-1 target was absent. The interaction zone was defined as the rectangular area 8 cm extended from the sides of the wire-mesh enclosure. Stressed animals were classified as resilient if the social interaction ratio was greater than 1. Otherwise, they were classified as susceptible.

### Two-photon imaging

The laser-scanning two-photon microscope was equipped with a resonant scanner unit for imaging at a frame rate of 29.8 Hz (Movable Objective Microscope, Sutter Instrument). The excitation source was a Ti:Sapphire laser tuned to a wavelength of 920 nm (Chameleon Ultra II, Coherent). The excitation was focused onto the brain by a high-numerical aperture water immersion microscope objective (XLUMPFLLN20X/1.0, Olympus). The microscope was controlled by a computer through the ScanImage software (Vidrio Technologies). Fluorescence was collected in two channels with GaAsP photomultipler tubes behind bandpass filters from 500 – 550 nm for GCaMP6 and from 575– 645 nm for mRuby2. At the beginning and end of each imaging session, additional images were collected with an excitation wavelength of 1040 nm for mRuby2 to ensure the field of view remains the same.

### Analysis of behavioral data

Time stamps for tongue licks and fluid deliveries were logged by Presentation in a delimited text file. The text files were parsed for analyses in MATLAB (MathWorks, Inc.). Licks that were separated in time by less than 0.5 s were considered to be part of a bout, also referred to as a lick cluster (29). Consummatory licks were defined as those licks within a bout that occurred after the reinforcement. When counting the number of rewards earned per block, we included the first reward that was needed to initiate the 60-s block. For all of the analyses, sessions in which animals initiated less than 4 blocks were excluded. That is, to be included in the analyses, they had to complete at least one cycle of 0%, 3%pc, 10%, and then 3%nc blocks. Moreover, for longitudinal analyses involving repeated social defeat, we only included animals in which we had at least one session of behavioral data for each of the pre-stress (day -1 and 1), during-stress (day 5, 7, and 9), and post-stress (day 11, and 13) stages.

### Analysis of behavioral data – modeling

The motivation and rationale behind the behavioral model had been described in detail in previous studies (34, 35). Our formulation was identical except the addition of marginal utility to capture satiety. Briefly, the equation was:

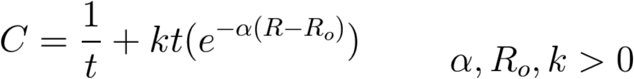

where *C* was the total cost, *t* was the latency between successive action, *k* was the reward rate, *α* was the rate for an exponential utility function, and *R*_*o*_ was the reward change-point when the utility would start to decline. For an exponential utility function, the marginal utility should have a scaling factor, but we did not include it because the factor was effectively absorbed into *k*. The reward rate *k* depended on the reinforcement type:

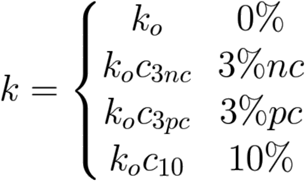

where *k*_*o*_ was the reward rate for water reinforcement, and *c*_*3nc*_, *c*_*3pc*_, *c*_*10*_ represented scaling factors to account for the different reward rates for the other types of reinforcements. The goal of the animal was to minimize the total cost. The optimal latency T to achieve the goal could be found by setting the first derivative to zero:

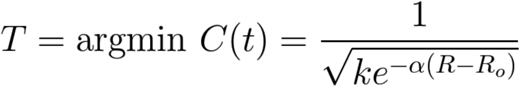

The equation was fitted separately for each session of behavioral data. There were 5 free parameters: *k*_*o*_, *c*_*3nc*_, *c*_*3pc*_, *c*_*10*_, *R*_*o*_. The parameter *α* was fixed at 0.7733, which was the median value determined by model fitting using calibration data (performance in a task in which mice perform the same FR10-FI5 tandem schedule to obtain only water rewards). For each fit, multiple initial conditions were tested, involving a range of parameters: *k*_*o*_ = 10-4, 10-3, 10-2; [*c*_*3nc*_ *c*_*3pc*_ *c*_*10*_] = [1 1 1], [1 1 2], [1 2 4], [1 2 8], and *R*_*o*_ = 100, 300, 500, corresponding to a total of 24 initial conditions. The best fit would minimize the sum of squared error, where error was defined as the difference between the predicted and measured number of earned rewards for each block. The predicted number of earned reward per block was simply the 60-s duration divided by the optimal latency, plus one to include the first earned reward that initiated the block.

We tested a number of modifications to the models to make sure that it is not sensitive to specific assumptions. One, for marginal utility, we replaced the exponential utility function with an isoelastic utility function. This modification typically yielded larger errors for the fits. Two, instead of fixing the *α* parameter, we treated it as another free parameter. This increased the number of initial conditions and therefore computation time, even though we found that the exact value of *α* tended to have minimal influence on the sum of squared error. We fitted *α* using a small number of sessions, and used the median as the fixed value, and then fit the entire data set. Three, another definition for error would be to minimize the sum of extra cost incurred for each action. Extra cost would be the difference between cost from ideal latency and cost from actual measured latency. This modification greatly increased the computation time. When tested on a dozen of sessions, this generated similar results as our primary approach.

### Analysis of neural data

Regions of interest corresponding to cell bodies were selected manually using a custom MATLAB graphical user interface. The regions of interest from the first imaging session was then also used for subsequent imaging sessions, sometimes with the locations translated slightly to match the actual image. For each cell body, the pixel values within the region of interest were summed to generate the time-lapse fluorescence signal, *F*(*t*). The fractional change of fluorescence was *dF*/*F*(*t*) = (*F*(*t*) – *F*_*o*_(*t*)) / *F*_*o*_(*t*) where *F*_*o*_(*t*) was the baseline fluorescence determined by finding the 10^th^ percentile of *F*(*t*) values within a 10-minute moving window.

To characterize spontaneous activity, for each cell, the fluorescence transients *dF*/*F*(*t*) was converted into inferred events using a peeling algorithm (52, 53). The peeling algorithm worked by template matching, and had been extensively demonstrated to be reliable for a wide range of signal-to-noise ratios, imaging speeds, and firing rates in numerical simulations (53). For all of the analyses, we set parameters to run the peeling algorithm (A1: 30; tau1: 1; frameRate: 30; with the rest set to default values). The inferred event rate was the number of inferred events divided by the duration of the no-lick interval. Cells were excluded from further analyses if they were inactive (overall rate of inferred event < 0.02 Hz) or had low signal-to-noise ([95 percentile of *dF*/*F*(*t*) – 50 percentile of *dF*/*F*(*t*)] < 1.2). These two criteria were determined by initially plotting the overall rate of inferred event against the signal-to-noise for all cells and identifying thresholds such that each cell would have distinguishable values in terms of these two metrics. We suspect that ‘cells’ that do not meet these criteria reflect ROIs that have drifted axially over time and therefore no longer contain meaningful fluorescence transient. We aimed to be conservative with these criteria such that only cellular fluorescence transients are included in the analysis. In total, we analyzed 1832 ROIs across all imaging sessions. The number of ROIs meeting the criterion for inferred event rate is 1319. The number of ROIs meeting the criterion for signal-to-noise is 1101. The number of ROIs meeting both criteria is 1062. Across days, any gap in measurements would be filled using linear interpolation. For example, if a cell was tracked on day -1, 1, 3, 7, 9, and 11, but missing day 5 and 13, then the value for day 5 would be interpolated using values from day 3 and 7, whereas day 13 would remain blank because it could not be interpolated. Additionally, for analyses of long-term activity changes, cells were excluded if they could not be tracked for at least the sessions on day -1 and day 1. For analyses of long-term activity changes, cells were excluded if they could not be tracked for at least 5 sessions.

For hierarchical clustering, we used cells from all animals. For each cell, we used the inferred event rates for the first 7 sessions (i.e., days -1, 1, 3, 5, 7, 9, and 11), and calculated the normalized change from baseline relative to the inferred event rate in day -1. The normalization was the difference divided by the sum, such that the normalized activity change was bounded by -1 and 1. We applied hierarchical clustering, based on Euclidean pairwise distance and agglomerating based on unweighted average distance between clusters, with a pre-determined cluster number of 5. We tested how varying the parameters would influence the clustering analysis. In particular, the analysis yielded similar conclusions based on correlation pairwise distance, and agglomeration linkage based on centroid distance or ward (inner squared distance). For other pairwise and cluster distance measures, the procedures would generate clusters that were not clearly separated, which was obvious by visually inspecting the pairwise similarity matrix. We tested changing the pre-determined cluster number from 3 to 10 with a step size of 1. Increasing the number beyond 4 did not yield any major division of a cluster, but rather led to more very small clusters containing only a few cells.

## Data availability

The behavioral and imaging data, as well as MATLAB code, are available on GitHub [link to be added].

### Acknowledgements

We thank Ian Stevenson for providing code to generate the beeswarm plot. This work was supported by National Institute of Mental Health grants R01MH112750 (A.C.K.) and R21MH110712 (A.C.K.), NARSAD Young Investigator Grant (A.C.K.), NARSAD Young Investigator Grant (F.B.), HHMI Medical Research Fellowship (M.H.), National Institutes of Health training grant T32NS041228 (M.J.S.), and National Science Foundation Graduate Research Fellowship DGE-1122492 (M.J.S.).

## Author contributions

F.B., M.H., and A.C.K. designed the research. F.B. and M.H. performed the behavioral studies. F.B. the imaging experiments. M.J.S. and F.A. provided code for analyzing imaging data. Y.S. and M.R.P. assisted with the social defeat paradigm. F.B., M.H., and A.C.K. analyzed the data. F.B. and A.C.K. wrote the paper, with input from all other authors.

## Competing interests

The authors declare no competing interests.

